# Unveiling Humoral and Cellular Immune Responses to SARS-CoV-2 in Head and Neck Cancer: A Comparative Study of Vaccination and Natural Infection in Romania

**DOI:** 10.1101/2024.10.27.620493

**Authors:** Luminita Mărutescu, Alexandru Enea, Nefeli-Maria Antoniadis, Marian Neculae, Diana Antonia Costea, Marcela Popa, Elena Dragu, Elena Codrici, Violeta Ristoiu, Bianca Galateanu, Ariana Hudita, Gratiela Gradisteanu Pircalabioru, Abdelali Filali-Mouhim, Veronica Lazăr, Carmen Chifiriuc, Raluca Grigore, Petronela Ancuta

**Author notes:** **Email addresses of the authors:**,;,;, and. **Lead contact: Petronela Ancuta**. equal contribution (Luminita Mărutescu, Alexandru Enea). **Corresponding authors mailing addresses**: **Petronela Ancuta**, Ph.D., CHUM-Research Centre, 900 rue Saint-Denis, Tour Viger R, room, R09.416, Montreal, Quebec H2X 0A9, Canada, extension #35744; **Raluca Grigore**, M.D., Ph.D., University of Medicine and Pharmacy “Carol Davila” Bucharest, Dionisie Lupu 37, Bucharest, 4192910, Romania; **Carmen Chifiriuc**, Ph.D.: University of Bucharest, Lab. of Microbiology-Immunology, Aleea Portocalelor no. 1-3, Bucharest 060101, sect 6, Romania; **Veronica Lazar**, Ph.D., University of Bucharest, Lab. of Microbiology-Immunology, Aleea Portocalelor no. 1-3, Bucharest 060101, sect 6, Romania.

## Abstract

**Background:** To fill the knowledge gap regarding the antiviral immunity in oncologic patients, we performed a comparative study on natural/vaccine-induced SARS-CoV-2 immunity in head and neck cancer (HNC) in Romania.

**Methods:** Blood was collected from HNC (n=49) and controls (n=14), stratified as vaccinated (RNA/adenovirus-based vaccines), convalescent, and hybrid immunity. Plasma IgG/IgA antibodies (Abs) against Spike (S1/S2), receptor binding domain (RBD), and nucleocapsid (NC), and cytokines were quantified using the MILLIPLEX® technology. The frequency/phenotype/isotype of RBD-specific B-cells were studied by flow cytometry using tetramers (Tet^++^). Cell proliferation in response to Spike/NC peptides was monitored by carboxyfluorescein succinimidyl ester (CFSE) assay. A longitudinal follow-up was performed on n=25 HNC.

**Findings:** Levels of S1/S2/RBD-specific IgG/IgA Abs were similarly high in HNC and controls, but significantly increased in convalescent/hybrid *versus* vaccinated HNC. NC-specific IgG/IgA Abs were only detected in convalescent/hybrid immunity groups. The frequency of Tet^++^ B-cells in HNC was similar to controls, irrespective of the immunization status, and correlated positively with RBD IgG/IgA Abs and negatively with the time since immunization (TSI). Compared to total B-cells, Tet^++^ were enriched in CD27^+^ memory phenotype and IgG/IgA isotype. A linear regression model identified Spike S2 IgG and NC IgA Abs as strong positive predictors of Tet^++^ frequencies, while IL-6 was a marginally significant negative predictor. Tet^++^ frequency remained stable at median TSI of 341 *versus* 117 days, despite a decline in memory phenotype.

**Interpretation:** HNC participants mount efficient and durable SARS-CoV-2 humoral immunity, with RBD-specific IgG/IgA Abs and Tet^++^ B-cells representing the major immunization outcomes.

## INTRODUCTION

The severe acute respiratory syndrome coronavirus 2 (SARS-CoV-2)^1^, a positive sense single-stranded RNA virus from the *Coronaviridae* family^2–4^, emerged in December 2019, in the Wuhan region of China^5,6^, and represents the etiological agent of the novel coronavirus disease-2019 (COVID-19)^4,7–9^. As of October 2023, the Centre for Systems Science and Engineering at Johns Hopkins University (https://coronavirus.jhu.edu/map.html) reported that SARS-CoV-2 has infected over 676 million people and resulted in over 6.8 million reported deaths, with over 13 billion doses of SARS-CoV-2 vaccines administered worldwide.

The existent knowledge on SARS-CoV-1, another coronavirus that emerged in 2002 in China^10,11^, facilitated the rapid discoveries on SARS-CoV-2^4^. Similar to SARS-CoV-1^12^, SARS-CoV-2 enters target cells (*e.g.,* epithelial cells in the respiratory tract) *via* the angiotensin converting enzyme 2 (ACE2), which serves as the primary entry receptor^13^. The secondary entry receptor, the transmembrane protease serine 2 (TMPRSS2), is essential for the Spike envelope glycoprotein cleavage^4,13–15^. Several other co-factors contribute to SARS-CoV-2 entry, including neuropilin-1 (NRP1)^16,17^ and CD147^18,19^. The ACE2 and TMPRSS2 are co-expressed at high levels in the nasal epithelium, while epithelial cells from the small airway mainly express TMPRSS2^20^. This differential expression of SARS-CoV-2 receptors in the upper *versus* lower respiratory tract points to anatomic site-specific particularities in the mechanisms of viral entry^15,21^. The pathogenicity of SARS-CoV-2 infection depends on its ability to make a transition from the nasal to the pulmonary compartment^22,23^. In the lungs, SARS-CoV-2 infection mediates a series of mechanisms leading to cilia dysfunctions^24,25^. The impaired mucociliary clearance during severe COVID-19 facilitates viral replication and dissemination into the pulmonary tissue^26^. Finally, the incapacity of the innate/adaptive immune system to contain SARS-CoV-2 infection at mucosal barriers leads to the translocation of viral RNA in the plasma, viremia being correlated with the severity of COVID-19 disease and morbidity^27^.

In the context of the COVID-19 pandemic, new generation mRNA-based vaccines were designed and used at large scale for SARS-CoV-2 vaccination worldwide, starting early 2021^15,28^. Other SARS-CoV-2 vaccines included adenovirus vector-based, inactivated, and recombinant protein vaccines^15,28,29^. Large-scale immune monitoring studies were performed worldwide^30^, proving the efficacy of SARS-CoV-2 vaccines to promote robust humoral/cellular immunity^31–36^. Tools were developed to distinguish between immunization induced by vaccination and natural infection, using the concomitant measurement of Spike [S1, S2, receptor binding domain (RBD) antibodies (Abs induced by vaccination and natural infection] and nucleocapsid (NC) Abs, induced only by natural infection^32,33,37,38^. Early studies demonstrated the superior ability of IgA compared to IgG Abs to neutralize the SARS-CoV-2 virions^39^ and supported the importance of IgA Abs for efficient immune protection at respiratory mucosal barriers^22,23,35^. This led to the most recent design of intranasal vaccines to induce strong local immune responses^40^.

SARS-CoV-2 vaccines protect against the risk for severe COVID-19 pathology and reduced the viral transmission at population level, although the goal of sterilizing immunity was not achieved^41–45^. The later limit is caused by the emergence of novel variants of concern (VOC; *e.g.,* Alpha, Beta, Delta, Omicron), which were functionally distinct compared to the ancestral D614G strain^15,46^, and exhibit the ability to escape Abs neutralization^47–49^. This forced the updating of vaccine composition to align with the contemporary VOCs^50–53^ and stressed the need for the design of new therapeutic strategies to treat severe SARS-CoV-2 infection using universal antivirals and immunotherapies^28,54,55^. New efforts are still needed to continue the fight against this emergent virus using efficient preventive and curative approaches^56^.

The severity of COVID-19 symptoms are associated with age, sex, metabolic deficiencies and comorbidities^9,57–62^, as well as host genetic factors^63–66^, including type I IFN gene polymorphism^65^. Both innate and adaptive immunity protect against SARS-CoV-2 disease severity^22,32,33^, with mucosal barrier innate immunity being essential to limit the viral spread from the upper respiratory tract into the lungs^67–69^. SARS-CoV-2 natural immunity and vaccine efficacy is compromised in high-risk groups, including multiple sclerosis^70,71^, kidney transplant^72^, HIV-1 infection^73^, and cancer^74–78^. Although cancer patients exhibit altered immune responses and are considered at risk for COVID-19^79–84^, the long-term humoral and cellular immune responses against SARS-CoV-2 have not been extensively studied in different types of malignancies^78,85^, such as head and neck cancer (HNC)^86^. In 2020, HNC was the 7th most common cancer worldwide^87^, predominant in males, with smoking being the main risk factor^88–91^.

To address this knowledge gap, we performed a detailed immune monitoring study to document the quality and duration of natural and vaccine-induced SARS-CoV-2 humoral and cellular immunity in a cohort of patients with HNC (n=49) admitted for oncologic treatment at Coltea Hospital, Bucharest, Romania. A control group without oncologic conditions (n=14) was recruited in parallel. To this aim, we used multiplex bead approaches for the simultaneous quantification in the plasma of 8 distinct types of SARS-CoV-2 Abs, i.e. Spike (S1, S2 and RBD) and nucleocapsid (NC) Abs of IgG and IgA isotypes, as well as 25 cytokines. Also, we used polychromatic flow cytometry on peripheral blood mononuclear cells (PBMCs) to explore *ex vivo* the frequency of SARS-CoV-2-specific B-cells, binding RBD tetramers (Tet^++^), and to measure the proliferation potential of B-cells and T-cells in response to SARS-CoV-2 Spike and NC proteins. A linear regression model identified Spike S2 IgG and NC IgA Abs as strong immunological predictors of major immunity outcomes, here identified as the RBD Abs of IgG/IgA isotypes with neutralization potential and Tet^++^ B-cells as the source for these Abs. Finally, a longitudinal follow-up in a group of HNC participants (n=25) revealed the persistence of Tet^++^ B-cells up to 717 days post-immunization (median: 341 days). These findings offer important insights into the quality and duration of natural and vaccine-induced immunity to SARS-CoV-2 in patients with HNC. This knowledge could help shape health policies for the medical management of HNC patients.

## MATERIALS AND METHODS

### Ethics statement

This study using biological samples from human subjects was approved by the Institutional Review Board (IRB) of the University of Bucharest, Bucharest, Romania (Ethical approval:13/26.04.2021) and the IRB of the Col ea Hospital, Bucharest, Romania (Ethical approval: PV 21/16.06.2021). This study was conducted in accordance with the ethical principles for biomedical research involving human subjects established by the Declaration of Helsinki in 1975. Informed consent was obtained from all participants involved in the study. Written informed consent has been obtained from the patients to publish this paper. Blood collection methods were carried out in accordance with relevant guidelines and regulations.

### Study participants

The study participants, represented by patients with HNC (n=49) and the control group [CTR, n=14; without neoplasm history, with (n=7) or without (n=7) other otorhinolaryngology pathology] were recruited at the Coltea Hospital, Bucharest, Romania, between August 2021 and March 2022 (Supplemental Figure 1). The description of the study participants is detailed in Table 1 in terms of sex, age, body mass index (BMI), SARS-CoV-2 infection symptoms, immunization status (vaccination and/or natural infection), time since immunization (TSI), diabetes, and oncologic pathology details. Peripheral blood samples were collected from participants at one visit (for all participants) or two subsequent visits (n=25 HNC and n=3 CTR participants). Plasma and peripheral blood mononuclear cells (PBMC) were isolated from whole blood and stored at −80 °C until use.

**Table 1:**
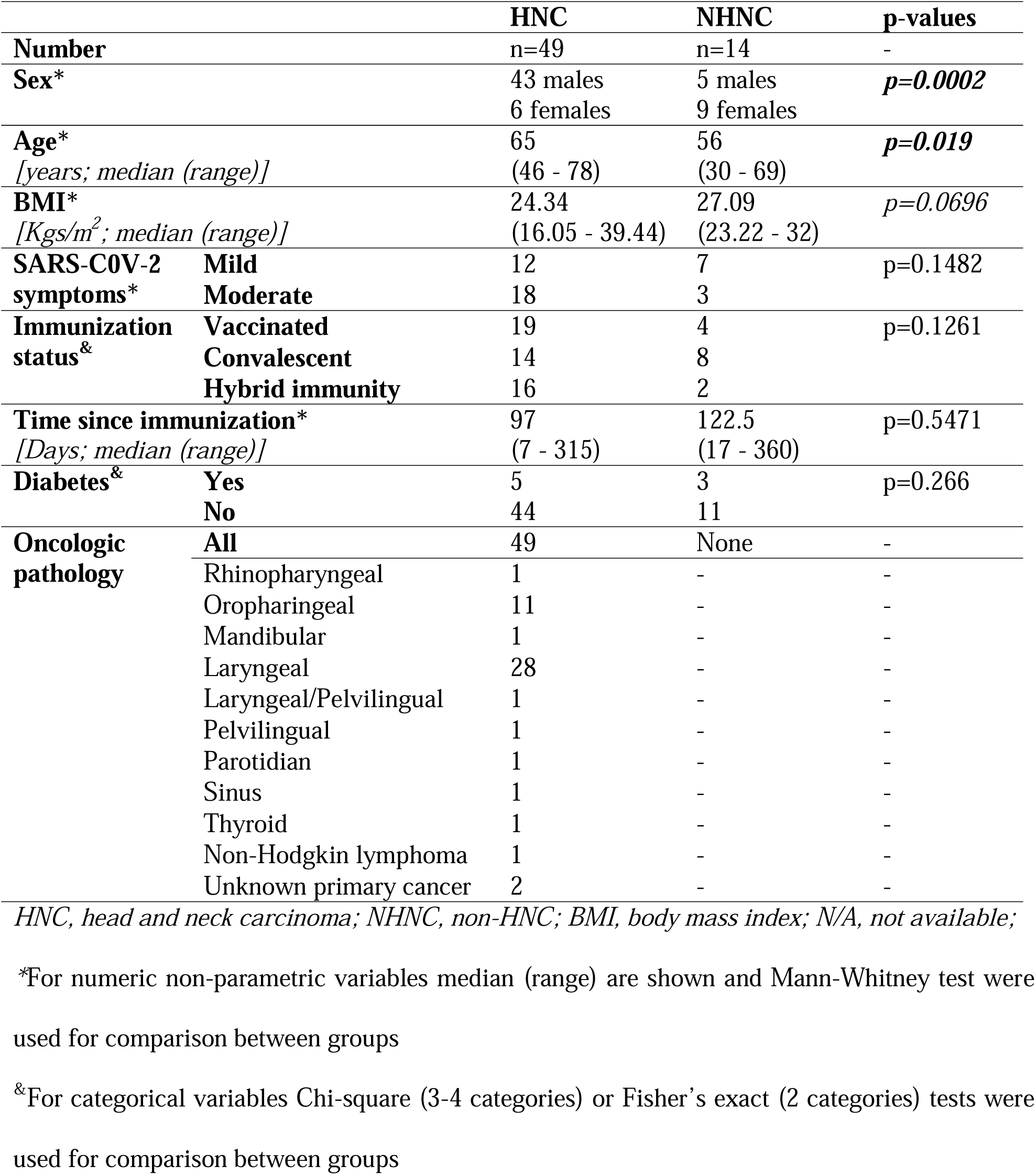
Description of the study cohort.

### Plasma and PBMC sample separation

Peripheral blood (10-40 ml blood per participant) was collected by venipuncture on EDTA tubes (BD vacutainer; Plymouth, UK), and processed within two hours. Blood centrifugation was performed at 2,070 rpm for 10 minutes at room temperature. The superior plasma fraction was collected, dispatched in 5-10 aliquots on 1 ml, and stored at −80°C until use. PBMCs were isolated from whole blood upon Ficoll–Paque (GE17-1440-02 Ficoll® Paque Plus; Uppsala, Sweden) density gradient centrifugation at 2070 rpm for 30 minutes at room temperature. PBMC were cryopreserved in fetal bovine serum (FBS, 2394341RP; Gibco, Mexico) containing 10% DMSO (D4540-1L; Sigma, Burlington, Massachusetts, USA) at −80°C, until further use, as we previously described^92,93^.

### Luminex detection of plasma IgG and IgA Antibodies to SARS-CoV-2

Specific IgG and IgA Abs profiles against four SARS-CoV-2 viral antigens [receptor binding domain (RBD), spike 1 subunit (S1) and spike 2 subunit (S2) of spike protein, and nucleocapsid (NC)] were monitored in the plasma samples using Millipore’s MILLIPLEX® MAP SARS-CoV-2 Antigen EMD panel IgG and IgA (Merck, Darmstadt, Germany), according with the manufacturer’s instructions. Briefly, 1/100 diluted plasma samples were incubated with fluorescent magnetic beads coated with the following viral recombinant SARS-CoV-2 proteins: Spike S1, Spike S2, RBD, and NC. Then, PE-conjugated anti-human IgG or PE-conjugated anti-human IgA were added to detect beads coated with plasma Abs. The antigen-Abs complexes on the beads were analyzed using a Luminex® 200™ system (Luminex, Luminex Corp., Austin, TX, USA) generating the median fluorescence intensity (MFI)) of the signal per 100 beads. Data acquisition and analysis were performed using xPONENT 4.2 software (Luminex, Luminex Corp., Austin, TX, USA); the calibration curves were generated with a 5-parameter logistic fit. Results were expressed as relative fluorescence units (RFU).

### Flow cytometry quantification of SARS-CoV-2 RBD-specific B-cells in blood

The detection of SARS-CoV-2-specific B-cells present in PBMC, was realized using the SARS-CoV-2 RBD B-cell MicroBead kit (Miltenyi Biotec, Germany), according to the manufacturer’s protocol. Briefly, 10^7^ PBMCs were stained with two solutions of RBD tetramers (Tet) conjugated with PE and PE-Vio770. Tet were generated by incubating biotin-labeled recombinant SARS-CoV-2 RBD protein with Streptavidin-PE and Streptavidin-PE-Vio770. Concomitant surface staining was performed with fluorochrome-conjugated Abs: CD19 APC-Vio770 Abs (clone LT19, isotype: mouse IgG1k) and CD27 VioBright FITC Abs (clone M-T271, isotype: mouse IgG1k); and/or with the anti-isotype Abs VioBlue-conjugated anti-IgG (clone IS11-3B2.2.3, isotype: mouse IgG1k), VioGreen-conjugated anti-IgA (clone IS11-8E10, isotype: mouse IgG1k), and APC-conjugated anti-IgM clone (PJ2-22H3, isotype: mouse IgG1). After incubation at room temperature for 30 minutes, the PBMCs were washed with PEB buffer (PBS, pH 7.2, 0.5% bovine serum albumin (#SLCF3210 Merck-Sigma, USA) and 2 mM EDTA (Merck-Sigma, USA)) and analyzed by flow cytometry using a 3-laser Gallios flow cytometer (Beckman Coulter, USA). Cell debris, doublets and dead cells were excluded from the analysis based on the size/granularity scatter signals. CD19^+^ B-cells with and without a memory (CD27^+^) phenotype were analyzed for the presence of RBD-specific B-cells identified using the double discrimination method of tetramer positive (Tet^++^) cells (PE/PE-Vio770). The IgM, IgG and IgA isotype of total and Tet^++^ B-cells was also analyzed. Fluorescence minus one (FMO) strategy was used to establish the positivity gates, as previously reported^94^.

### Carboxyfluorescein succinimidyl ester (CFSE) proliferation assay

The CFSE proliferation assay was performed, as we previously reported^95,96^, to monitor the SARS-CoV-2-specific proliferation of immune cells from PBMC of HNC and NHNC study participants. Briefly, PBMCs (2×10^6^ cells/ml) were stained with 0.5 μM CFSE (BioLegend, SanDiego, CA, USA; B377681) in PBS 1X (Gibco, Geel, Belgium; 18912-014), for 8 minutes at 37 °C, and protected from light. After washing, the cells were suspended in proliferation media (RPMI 1640, L-glutamine and Hepes (Gibco, NY, USA; A10491-01) supplemented with 1% penicillin/streptomicin (Gibco; NY, USA; 15140122) and 10% human serum (ZenBio, Durham/USA; HSER-ABP, SER052318). After inoculation at a density of 0.25×10^6^ cells/well in 96-well microtiter plates (TPP; Saint Louis, USA; 96F92696), at a final volume of 250 µl/well, cells were stimulated with SARS-CoV-2 peptide pools (PepTivator® SARS-CoV-2 Prot_N 130-126-699; SARS-CoV-2 Prot_S 130127953) (0.6 nmol/mL each) (Miltenyi Biotec; Bergisch Gladbach, Germany). After 6 days of incubation at 37°C, 5% CO2, the cells were harvested and labelled on the surface with the following monoclonal antibodies (Abs) from Biolegend (San Diego, USA): APC-conjugated anti-CD3 Abs (clone Mouse IgGak OKT3), PE-conjugated anti-CD4 (clone Mouse IgG1k RPA-T4), PerCP-conjugated anti-CD8 (clone Mouse IgG1k SK1), APC-conjugated anti-CD19 (clone HIB19), PE-conjugated anti-CD56 (clone HCD56). Following staining, cells were washed twice with FACS buffer (PBS 1X, 0.02% NaN3, 10% FBS) and analyzed using the BD Accuri C6 Plus (BD Biosciences; New Jersey, USA) flow cytometer. The data analysis was performed using the BD Accuri C6 software (BD Biosciences, USA).

### Luminex quantification of plasma Th17 cytokine profiles

Plasma cytokine/chemokine levels were measured using the Human Th17 Magnetic Bead Panel (TH17MAG-14kMILLIPLEX MAP, Merck; Darmstadt, Germany) allowing the simultaneous quantification of the following cytokines: IL-1β, IL-2, IL-4, IL-5, IL-6, IL-9, IL-10, IL-12p70, IL-13, IL-15, IL-17A, IL-17E/IL-25, IL-17F, IL-21, IL-22, IL-23, IL-27, IL-28A, IL-31, IL-33, GM-CSF, IFN-γ, MIP-3α/CCL20, TNF-α and TNF-β. The assay was performed according with the manufacturer’s instructions. Briefly, 25 µl plasma samples were incubated to fluorescent magnetic beads coated with Abs against cytokines. Then, PE-conjugated anti-cytokine Abs were added to detect beads coated with cytokines. The antigen-Abs complexes on the beads were analyzed using the Luminex® 200™ system (Luminex, Luminex Corp., Austin, TX, USA) generating the median fluorescence intensity (MFI) of the signal per 100 beads. The use of a standard curve for each cytokine allowed the expression of the results as pg/ml.

### Statistical analysis

For comparisons between groups and correlations, statistical analyses were performed using the Prism 7 (GraphPad, Inc., La Joya, CA, USA) software. Statistical tests applied and p-values are indicated on the graphs and explained in the figure legends.

#### Linear regression

To investigate the association between humoral and cellular immune response parameters that predict specific SARS-CoV-2 immunity outcomes [*i.e.,* SARS-CoV-2 RBD IgG and IgA Abs (Abs with neutralization potential) and the frequencies of SARS-CoV-2 Spike-specific B-cells (Tet^++^ B-cells)], a linear regression model was implemented with adjustments for numerical (*i.e.,* age, BMI, TSI) and/or categorical variables (*i.e.,* sex, smoking, alcohol, toxic environment, diabetes). Regression coefficient as well as the p-values for each immune parameter were calculated (Supplemental Tables 1-4). P-values adjustment for multiple testing hypothesis was performed according to the method of Benjamin and Hochberg^97^, which controls the false discovery rate (FDR) with adjusted p-value cut-offs of 0.05. Multivariate predictive modeling was performed using least absolute shrinkage and selection operator (LASSO)^98^, a regression analysis method well suited for predicting outcome in the presence of many predictors p (p>>N) that performs both variable selection and regularization in order to obtain a subset of predictors (model) that minimizes the prediction error using leave-one-out Cross Validation technique. All statistical analyses were performed using the statistical package R version 4.2.3.

### Roles of funders

The funding institutions played no role in the design, collection, analysis, and interpretation of data. The funders played no role in data analysis and results interpretation, nor the writing of the manuscript.

## RESULTS

### Study participants

Study participants, including people HNC patients (n=49) and control individuals without cancer diagnosis (NHNC, n=14), were recruited during the period August 2021 and March 2022 (Supplemental Figure 1A). Table 1 depicts the description of the study cohort. The HNC and control groups differed in terms of age (median 65 and 56 years old, respectively; Mann-Whitney p=0.019) and male/female ratios (Fisher’s exact test p=0.0002), with HNC including more males than females and being older compared to controls. The HNC *versus* control groups were similar in terms of immunization status (*i.e.,* vaccinated, convalescent, hybrid immunity) and symptoms (i.e., mild to moderate, Fisher’s test) upon natural SARS-CoV-2 infection, as well as TSI, and diabetes prevalence. There was a tendency for BMI in HNC compared to NHNC (Mann-Whitney p=0.0696) (Table 1). Vaccinated HNC participants (20/49) received BioNTech/Pfizer BNT162b2 (n=10), Johnson & Johnson/Janssen Ad26.COV.2.S (n=4), Moderna mRNA-1273 (n=4), or Oxford–AstraZeneca vaccine (ChAdOx1 CoV-19) (n=1), while all the vaccinated controls (6/14) received the BioNTech/Pfizer BNT162b2 vaccine (<u>Supplemental File 1</u>). The oncologic pathology in the HNC group included rhinopharyngeal, oropharyngeal, mandibular, laryngeal, pelvilingual, parotidian, sinus, thyroid neoplasms, as well as participants with Non-Hodkin lymphoma and unknown primary cancers (Table 1; <u>Supplemental File 1</u>).

Within the HNC group, there were no statistical differences between males (n=43) and females (n=6) in terms of age, BMI and TSI (Supplemental Figure 2A-B). Within the HNC group there were no statistical differences in terms of age, BMI, and TSI between vaccinated, convalescent, and hybrid immunity participants (Supplemental Figure 2C).

### Humoral Immunity

#### Plasma SARS-CoV-2 IgG and IgA Abs

The analysis of plasma samples collected at the first visit (median TSI 107 days for HNC and 65 days for controls) showed detectable levels of SARS-CoV-2 IgG and IgA Abs (S1, S2, RBD, and NC) in the majority of participant samples (Figure 1), as late as 315 days post-immunization (<u>Supplemental File 1</u>). As expected, NC IgG and IgA Abs were detected only in convalescent and hybrid immunity HNC participants (Figure 1B and D).

**Figure 1.**
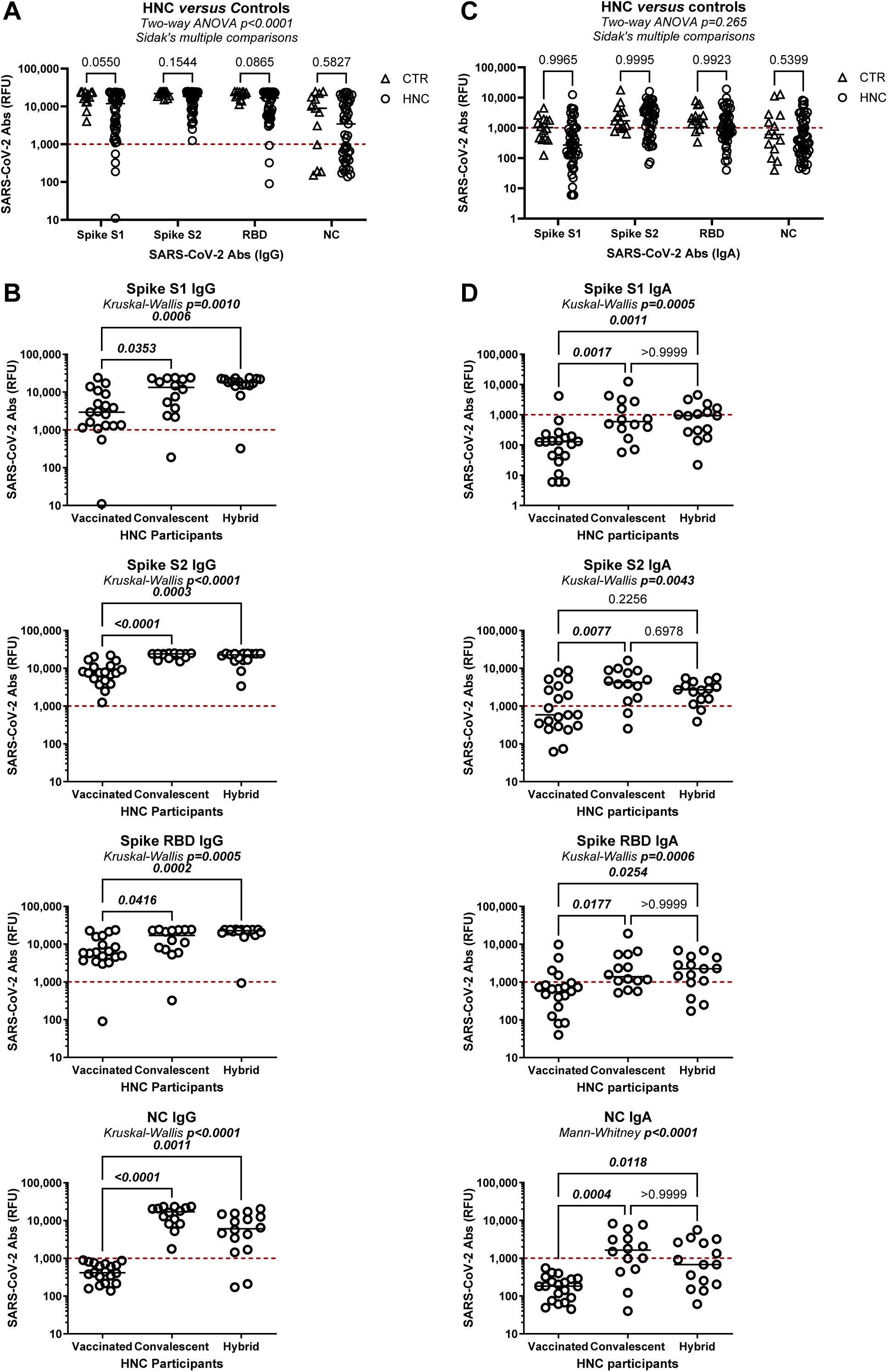
Plasma levels of SARS-CoV-2 IgG and IgA Abs in HNC participants upon vaccination and/or natural infection. Plasma isolated from whole blood served for the quantification of IgG **(A-B)** and IgA **(C-D)** Abs) against SARS-CoV-2 RBD, S1, S2 and NC using the Luminex xMAP-based multiplex assay. Results are expressed as Relative Fluorescence Units (RFU). **(A and C)** Shown are levels of SARS-CoV-2 RBD, S1, S2 and NC Abs of IgG (**A**) and IgA **(C)** isotype in the plasma of people with HNC (n=49) *versus* healthy controls (CTR; n=14). Two-way ANOVA test followed by Sidak’s multiple comparisons determined differences between groups. **(B and D)** Shown are plasma levels of SARS-CoV-2 RBD, S1, S2 and NC Abs of IgG (**B**) and IgA **(D)** isotypes among HNC participants stratified based on the immunization status as vaccinated, convalescents and hybrid immunity. Statistical significance was determined through the Kruskal Wallis test and subsequent Dunn’s multiple comparisons, with p-values indicated on the graphs.

Concerning the SARS-CoV-2 Abs of IgG isotype, HNC and control participants exhibited similar levels of plasma S1 and NC IgG Abs, while S1 and RBD IgG Abs titers tended to be lower in HNC *versus* controls participants (Figure 1A). Among HNC participants, S1, S2, and RBD IgG Abs levels were significantly higher in convalescent and hybrid immunity compared to vaccinated participants (Figure 1B). Spearman models revealed no statistically significant correlations between S1, S2, RBD, or NC IgG Abs titers and age, TSI and BMI (<u>data not shown</u>). Concerning SARS-CoV-2 Abs of IgA isotypes, their levels were overall lower when compared to IgG Abs levels, but statistically similar between HNC and controls (Figures 1A and 1C). Among HNC participants, the convalescent *versus* vaccinated status was associated with significantly higher levels of S1, S2, and RBD IgA Abs, with no statistically significant differences between convalescent and hybrid immunity status (Figure 2B). While age was not a predictor of IgA Abs levels, Spearman correlation models identified TSI as a negative predictor of plasma NC (p=0.0356, r=-0.3009), but not S1, S2, and RBD IgA Abs titers (<u>data not shown</u>). Finally, BMI positively correlated with RBD IgA Abs (p=0.0391; r=3121), but not with S1, S2 and NC IgA Abs (<u>data not shown</u>).

**Figure 2:**
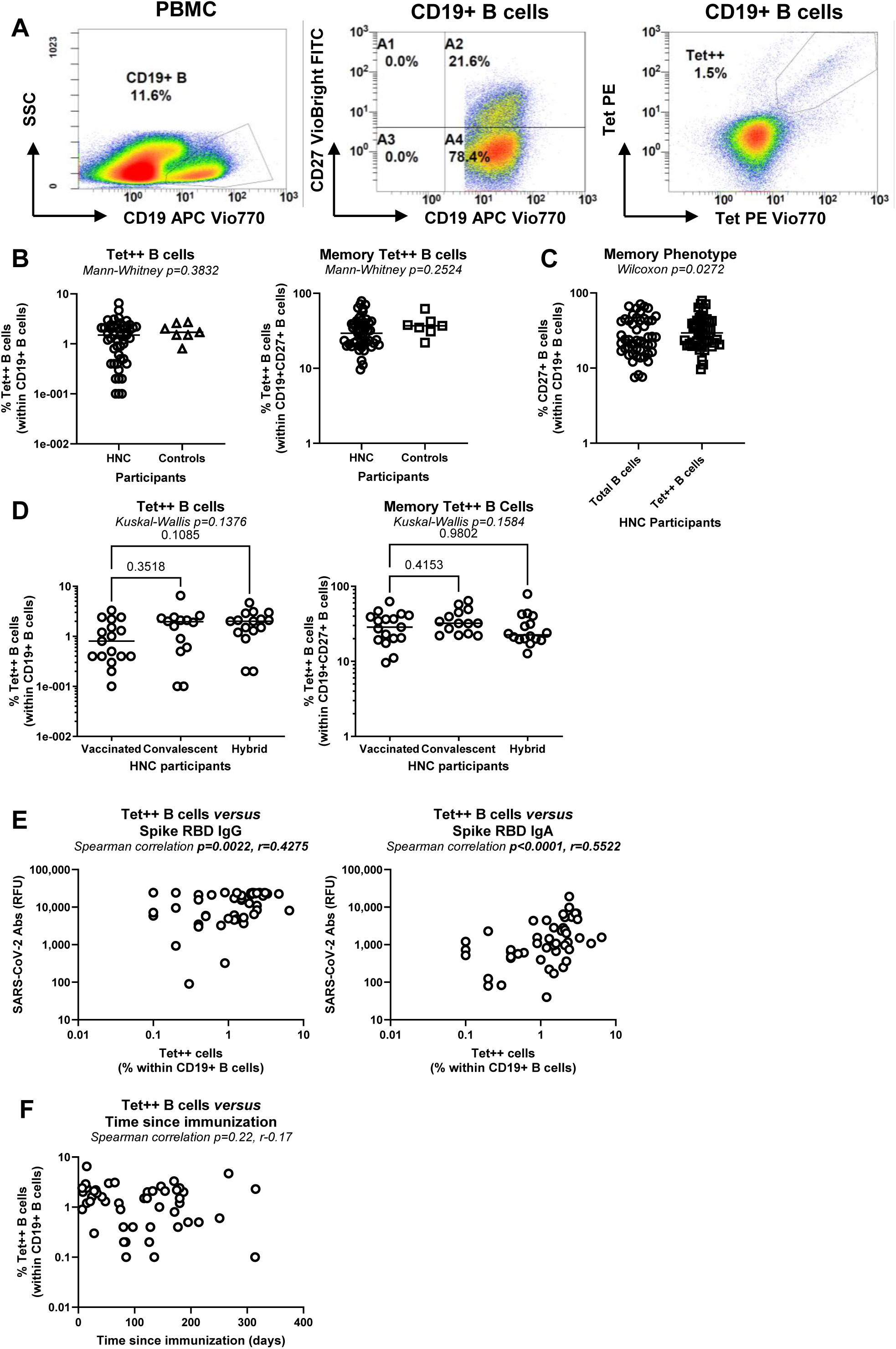
Frequency of RBD-specific B-cells in the blood of HNC participants upon vaccination and/or natural infection. PBMC were isolated from whole blood by Ficoll gradient density centrifugation. (**A**) Shown is the flow cytometry gating strategy in one representative control participant for the identification a phenotypic characterization of SARS-CoV-2 RBD-specific B-cells within PBMC *ex vivo*. Tetramers (Tet) were generated by incubating Biotin-labeled SARS-CoV-2 RBD-Protein with Streptavidin-PE and Streptavidin-PE-Vio770. Surface staining was performed concomitantly with CD19 and CD27 Abs conjugated with the APC Vio770 and FITC fluorochromes, respectively. Lymphocytes were identified based on their size and granularity as indicated by the forward scatter (FSC) and sideward scatter (SSC) signals. B-cells were identified by expression of CD19. Memory B-cells were identified as CD19^+^CD27^+^ cells. SARS-CoV-2 RBD-specific B-cells were identified as cells co-expressing Tet PE and Tet PE-Vio770 (Tet^++^). (**B-D**) Shown is the frequency of total (**left panel**) and memory (**right panel**) Tet^++^ B-cells in HNC *versus* control participants (**B**), the expression of CD27 on total *versus* Tet^++^ B-cells from HNC participants (**C**), and among HNC participants stratified based on the immunization status in vaccinated, convalescent and hybrid immunity (**D**). Mann-Whitney (**B**), Wilcoxon (**C**), and Kuskal-Wallis with Dunn’s multiple comparisons (**D**) determined differences between the two groups. (**E**) Shown are the Spearman correlation p and r values for the correlation between the frequency of total Tet^++^ B-cells within PBMC and plasma levels of SARS-CoV-2 RBD IgG (**E, left panel**) and IgA (**E, right panel**) Abs in HNC participants. **(F)** Shown is the correlation between the frequency of Tet^++^ B-cells and the time since immunization, with Spearman correlation p and r values indicated on the graphs.

Together, these results reveal the induction of robust SARS-CoV-2 IgG and IgA Abs responses in HNC participants upon vaccination and/or natural infection and point to the RBD-specific IgG and IgA Abs, which likely exhibit neutralization potential, as major outcomes of SARS-CoV-2 humoral immunity.

#### Circulating SARS-CoV-2-specific B-cells in HNC participants

To further characterize the SARS-CoV-2 humoral immunity, total B-cells (CD19^+^) where analyzed for their SARS-CoV-2 RBD specificity and memory phenotype (CD27^+^) by flow cytometry upon staining with SARS-CoV-2 RBD tetramers (Tet^++^) and CD27 Abs, respectively, using the gating strategy depicted in Figure 2A. Of note, the total (CD19^+^) and memory (CD19^+^CD27^+^) Tet^++^ B-cells were present at various frequencies in HNC participants, with no statistically significant differences when compared to control participants (Figure 2B). Within the HNC group, Tet++ B-cells were slightly enriched in the memory phenotype compared to total B-cells (median 29.3% *versus* 23.8% CD27^+^ B-cells, p=0.0272; Figure 2C), with not statistical differences in the frequency of total and memory Tet^++^ B-cells among the vaccinated, convalescent and hybrid immunity groups (Figure 2D). It is noteworthy that the frequency of Tet^++^ B-cells positively correlated with plasma levels of SARS-CoV-2 RBD Abs of IgG and IgA isotypes (Figure 2E). Of note, the % of Tet++ B-cells did not correlate with the TSI (Figure 2F), indicative of the stability of SARS-CoV-2 B-cell memory after infection/vaccination. In parallel, we investigated the IgM, IgG, and IgA isotype of total *versus* Tet^++^ B-cells, using the gating strategy depicted in Figure 3A. Among HNC participants, Tet^++^ B-cells distinguished from total B-cells by lower expression of IgM and higher expression of IgG and IgA (Figure 3B). This suggests a more effective immunoglobulin class switch from IgM to IgG or IgA following exposure to SARS-CoV-2. The isotype of Tet^++^ B-cells was similar in HNC and control participants (Figure 3C), as well as among HNC participants with different immunization status (*i.e.,* vaccinated, convalescent and hybrid immunity groups) (Figure 3D).

**Figure 3:**
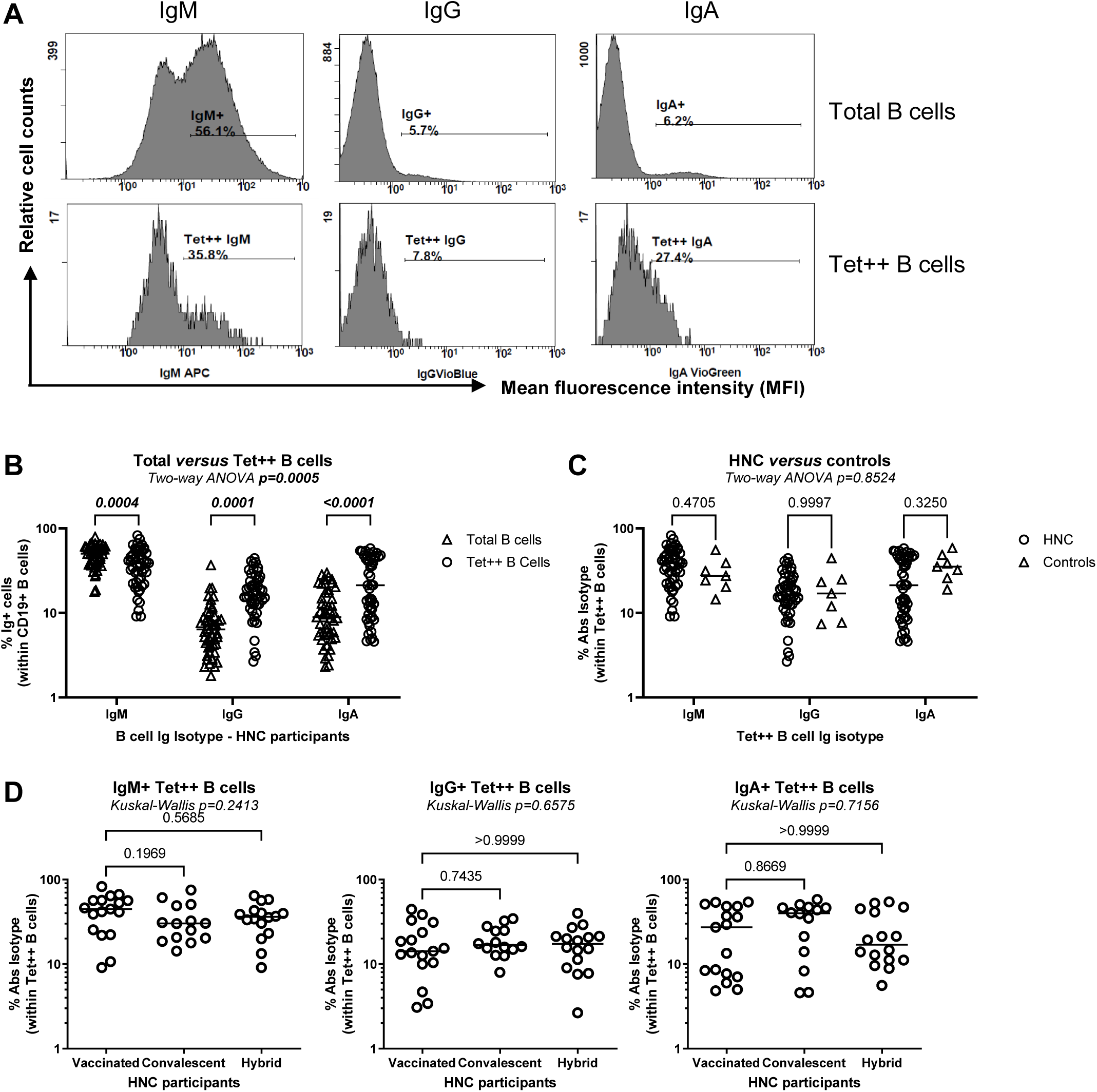
Isotypic characterization of RBD-specific B-cells in the blood of HNC participants upon vaccination and/or natural infection. (**A**) Shown is the gating strategy in one representative control participant for the isotypic characterization (IgM, IgG and IgA) of total and Tet^++^ B-cells, identified as shown in Figure 2A. (**B-D**) Shown are the isotype of total *versus* Tet^++^ B-cells within HNC participants (**B**), the isotype of Tet^++^ B-cells in HNC *versus* control participants (**C**), as well as the IgM (**D, left panel**), IgG (**D, middle panel**) and IgA (**D, right panel**) isotype of Tet^++^ B-cells in HNC participants classified based on their immunization status in vaccinated, convalescent and hybrid immunity (**D**). P-values for Two-way ANOVA followed by Sidak’s multiple comparisons (**B-C**) and Kruskall-Wallis and Dunn’s multiple comparisons (**D**) are indicated on the graphs.

Together these results revealed the presence of RBD-specific B-cells, with CD27+ memory phenotype and IgM/IgG/IgA isotypes that showed no significant differences between HNC participants and controls, irrespective of immunization status or TSI among the HNC group. Furthermore, the positive correlation between the frequency of RBD-specific B-cells and the plasma levels of RBD-specific IgG/IgA Abs support the idea that these circulating Tet^++^ B-cells are involved in producing neutralizing antibodies against SARS-CoV-2 and represent an important SARS-CoV-2 immunity outcome to monitor in immunity surveillance studies.

#### Proliferation of SARS-CoV-2-specific B-cells

In addition to the use of tetramers for the identification of SARS-CoV-2-specific B-cells *ex vivo*, the CFSE dilution assay was used in parallel to determine the proliferative capacity of B-cells producing SARS-CoV-2 Abs, as a read-out of their survival capacity *in vivo*. Results reveal the proliferation (CFSE^low^) of CD19^+^ B-cells upon cultivation of PBMC in the presence or the absence of recombinant SARS-CoV-2 Spike and NC proteins (Figure 4A-B). Statistical analysis shows a significantly higher proliferation of CD19^+^ B-cells upon exposure to SARS-CoV-2 Spike compared to NC protein (p=0.0007) (Figure 4C), consistent with the well described superior immunogenicity of Spike relative to NC^31–33^. The proliferation of B-cells in response to SARS-CoV-2 Spike but not NC showed differences among the HNC group, with a statistically significant increase for convalescent *versus* vaccinated participants (p=0.0052; Figure 4D).

**Figure 4:**
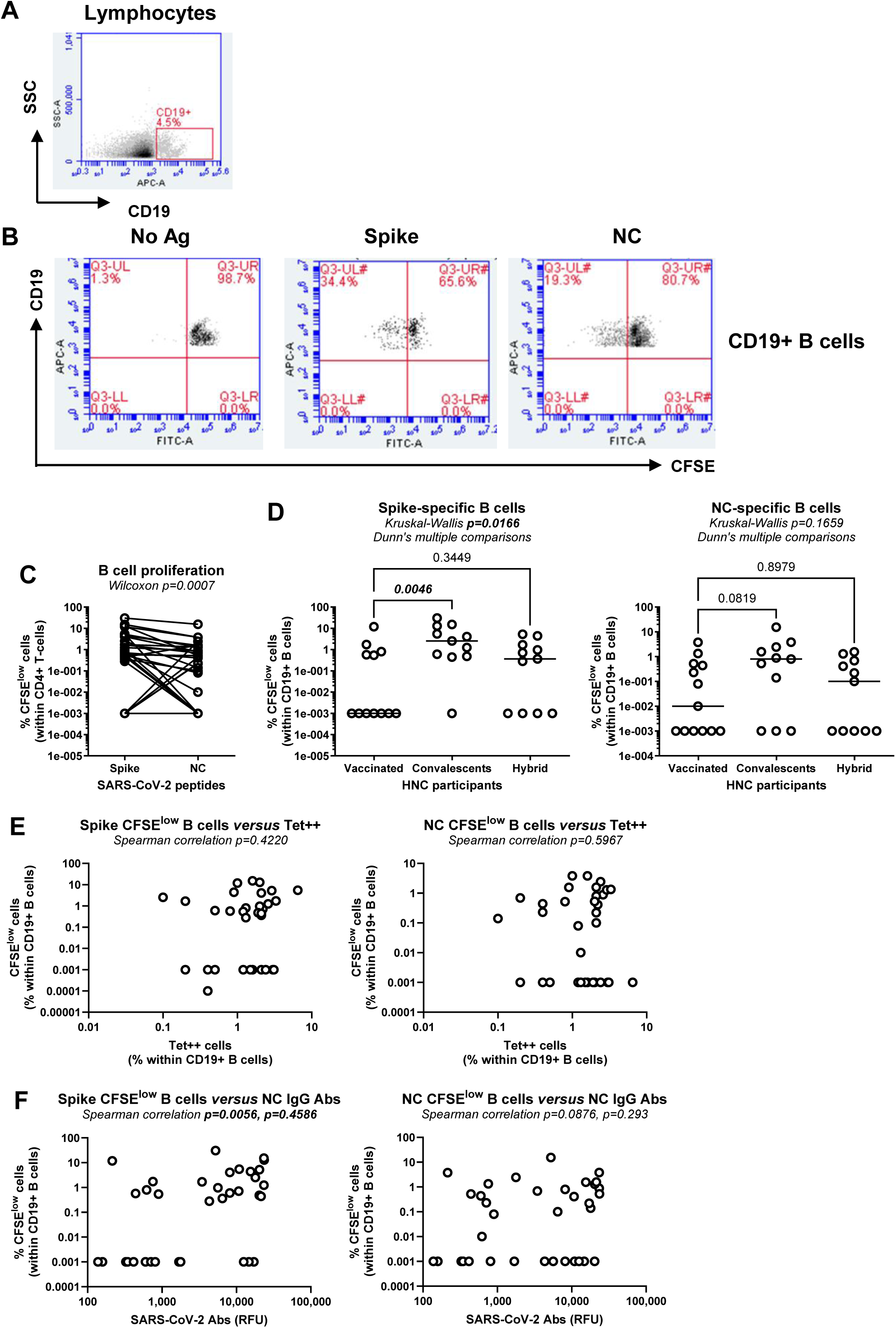
B-cell proliferation in response to SARS-CoV-2 Spike and NC proteins in HNC participants. PBMC were loaded with CFSE, exposed to recombinant SARS-CoV-2 Spike and Nucleocapsid (NC) recombinant proteins, and cultured for 6 days. Cells were harvested and stained with APC-conjugated CD19 Abs. The CFSE dilution in cells was used as an indicator of cell proliferation. Shown is the gating strategy used to identify CD19^+^ B-cells (**A**) and their proliferation (CFSE^low^) in response to SARS-CoV-2 Spike and NC antigens (Ag) compared to unstimulated control (No Ag) in one representative HNC participant (**B**). Numbers in the dot plots indicate the percentage of cells identified by manual gating. (**C**) Shown are differences in B-cell proliferation in response to SARS-CoV-2 Spike and NC in HNC participants (n=35), with the Wilcoxon test p-values indicated on the graph. (**D**) Shown is the proliferation of B-cells in response to SARS-CoV-2 Spike (**D, left panel**) and NC (**D, right panel**) Ags in HNC participants classified based on their immunization status in vaccinated, convalescent and hybrid immunity. Kruskall-Wallis and Dunn’s multiple comparison p-values are indicated on the graphs. (**E**) Shown are the correlations between the frequency of Tet++ B-cells in the blood and B-cell proliferation (CFSE^low^) in response to SARS-CoV-2 Spike (**E, left panel**) and NC (**E, right panel**) Ags. Spearman correlation p and r-values are indicated on the graphs.

To identify the predictors of SARS-CoV-2-specific B-cell proliferation, the Spearman correlation model was applied to analyze the relationship between the frequency/isotype of Tet^++^ B-cells, along with the plasma levels of SARS-CoV-2 Abs in HNC participants. The frequency of CD19^+^ B-cells proliferating in response to SARS-CoV-2 Spike and NC proteins did not correlate with the frequency of Tet^++^ B-cells (Figure 4E), suggesting that some Tet^++^ B-cells may have impaired survival capacity. Nevertheless, consistent with the predominant proliferation of B-cells in convalescent HNC participants (Figure 4D), plasma levels of SARS-CoV-2 NC IgG Abs positively correlated with the frequency of proliferating CD19^+^ B-cells, with a significant correlation for Spike-specific B-cells (Figure 4F, <u>left panel</u>) and a marginally significant correlation for NC-specific B-cells (Figure 4F, <u>right panel</u>).

Thus, B-cell proliferation in response to SARS-CoV-2 peptides was more robust in convalescent *versus* vaccinated HNC participants, being positively correlated with plasma levels of NC-specific IgG Abs.

### Cellular immunity

#### Proliferation of SARS-CoV-2-specific CD4+ and CD8+ T-cells

The contribution of CD4^+^ and CD8^+^ T-cells to SARS-CoV-2 immunity is well recognized in the general population^23^. In an effort to identify cellular components of SARS-CoV-2 antiviral responses among HNC participants, we investigated the proliferation of CD4^+^ and CD8^+^ T-cells in response to recombinant SARS-CoV-2 Spike and NC proteins (Figure 5A-B). Similar to results on B-cell proliferation (Figure 4C), SARS-CoV-2 Spike depicted superior immunogenicity compared to NC peptides for the induction of CD4^+^ T-cells (p=0.0011) and CD8^+^ T-cells (p=0.00167) proliferation (Figure 5C-D). The % of proliferating (CFSE^low^) CD4^+^ and CD8^+^ T-cells in response to SARS-CoV-2 Spike and NC peptides was similar in HNC participants regardless of the immunization status (Figure 5E-F). The frequency of CD4^+^ T-cells proliferating in response to SARS-CoV-2 Spike but not NC peptides positively correlated with the frequency of Tet++ B-cells (Figure 5G-H), consistent with the well documented cross-talk between CD4^+^ T-cells and B-cells in sustaining humoral responses^99^. Such correlations were not observed between proliferating CD8^+^ T-cells and Tet^++^ B-cells (<u>data not shown</u>).

**Figure 5:**
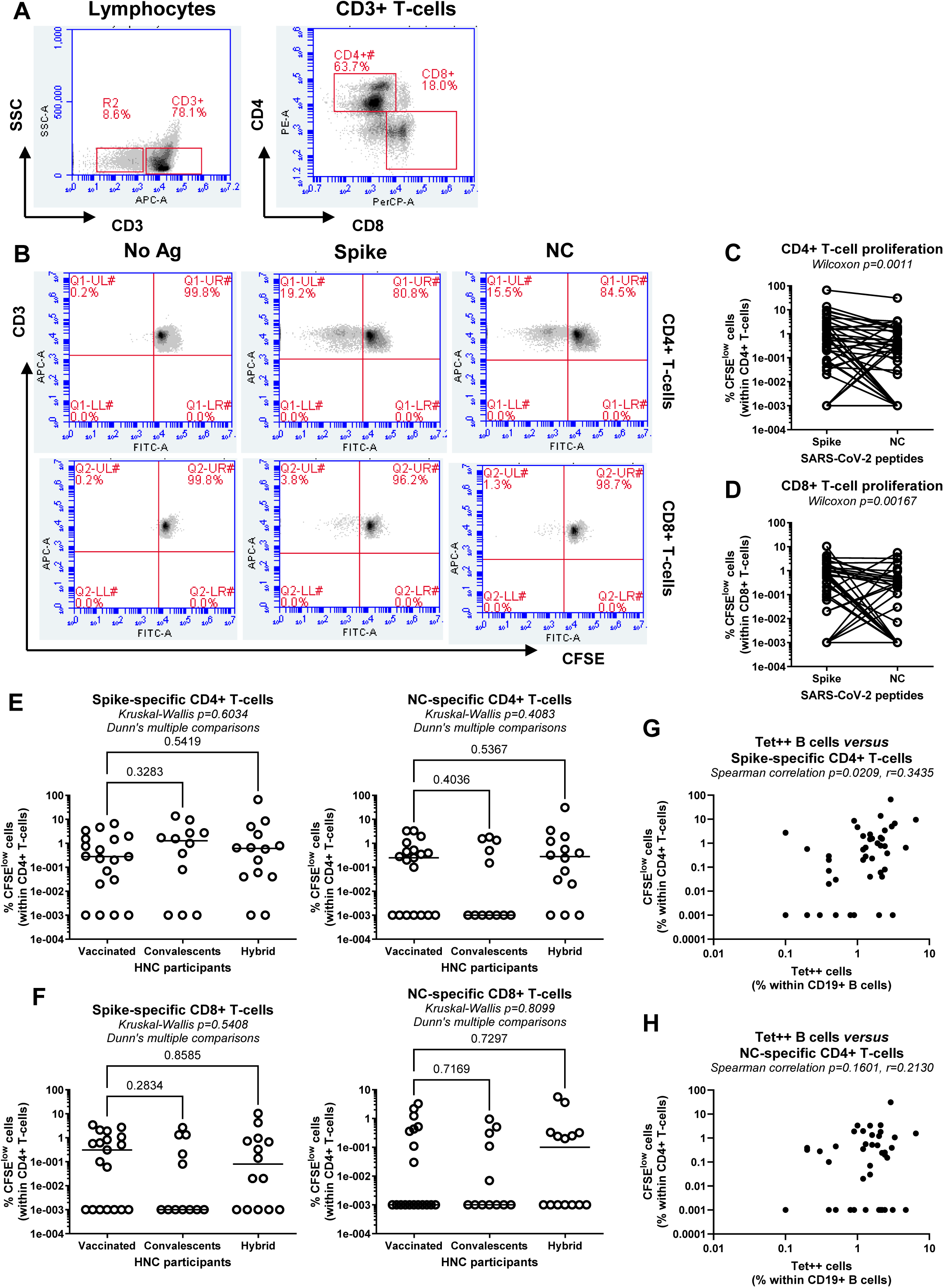
CD4^+^ and CD8^+^ T-cell proliferation in response to SARS-CoV-2 Spike and NC proteins in HNC participants. PBMC were loaded in CFSE, exposed to SARS-CoV-2 Spike and NC recombinant proteins and cultured, as described in Figure 4 <u>legend</u>. Cells were harvested and stained with CD3, CD4 and CD8 Abs conjugated with APC, PE and PerCP fluorochromes, respectively. (**A-B**) Shown is the gating strategy on one representative participant for the identification on CD3^+^ T-cells expressing CD4 or CD8 (**A**) and the CFSE dilution in CD4^+^ T-cells (**B, upper dot plots**) and CD8^+^ T-cells (**B, lower dot plots**) proliferating in response to SARS-CoV-2 Spike and NC recombinant proteins, compared to the No Ag control (**B**). Numbers in the dot plots indicate the percentage of cells identified by manual gating. (**C-D**) Shown are differences in the frequencies of CD4^+^ T-cells (**C**) and CD8^+^ T-cells (**D**) proliferating in responses to SARS-CoV-2 Spike *versus* NC protein, with Wilcoxon test p-values indicated on the graphs. (**E-F**) Shown are differences in the frequencies of CD4^+^ T-cells (**E**) and CD8^+^ T-cells (**F**) proliferating in responses to SARS-CoV-2 Spike (**left panels**) and NC (**right panels**) recombinant proteins among HNC participants classified based on the immunization status in vaccinated, convalescents and hybrid, with Kruskall-Wallis and Dunn’s multiple comparisons p-values indicated on the graphs. (**G-H**) Shown is the correlation between the frequency of Tet++ B-cells and the frequency of CD4^+^ T-cells proliferating in response to SARS-CoV-2 Spike (**G**) and NC (**H**) recombinant proteins. Spearman correlation p and r-values are indicated on the graphs.

### Systemic cytokines levels

Plasma levels of pro-inflammatory cytokines were reported to be elevated in patients with severe COVID-19^100^. To evaluate the systemic immune activation in relationship with the obove monitored humoral and cellular SARS-CoV-2 immunity readouts, plasma level of 25 cytokines were quantified using the Luminex®xMAP® technology, using low (*i.e.,* IL-17F, GM-CSF, IL-10, IL-22, IL-4, IL-23, IL17E/IL-25, IL-27, IL-31, TNF-β, IL-28A) and (*i.e.,* IFN-γ, CCL20, IL-12p70, IL-13, IL-15, IL-17A, IL-9, IL-1β, IL-33, IL-2, IL-21, IL-5, IL-6, TNF-α) high sensitivity standard curves, proportional with the physiological concentrations of specific cytokines in human plasma (Supplemental Figure 3A-B). Cytokines present at detectable levels in the plasma were further grouped as Th1/Th2/Th17 lineage and inflammatory cytokines (Figure 6A-B). Plasma levels of these cytokines tended to be increased in HNC compared to control participants, with statistically significant differences being observed for CCL20 (p<0.0001) and IL-33 (p=0.0011), and marginally significant differences for IL-21 (p=0.0597) (Figure 6A-B). The stratification based on the immunization status did not reveal statistically significant differences between groups for these three cytokines (Figure 6C). Finally, a Spearman correlation model aimed to identify cytokines that could be used as predictors of the major immunity outcomes. Of particular importance, although plasma levels of IL-6 did not distinguish HNC from control participants (Figure 6B), among HNC participants IL-6 levels negatively correlated with plasma levels of RBD IgG (p=0.0193, r=-0.3333) and IgA (p=0.0252, r=-0.3196) Abs, as well as with the frequencies of Tet^++^ B-cells (p=0.0024, r=-04233) (Figure 6D).

**Figure 6:**
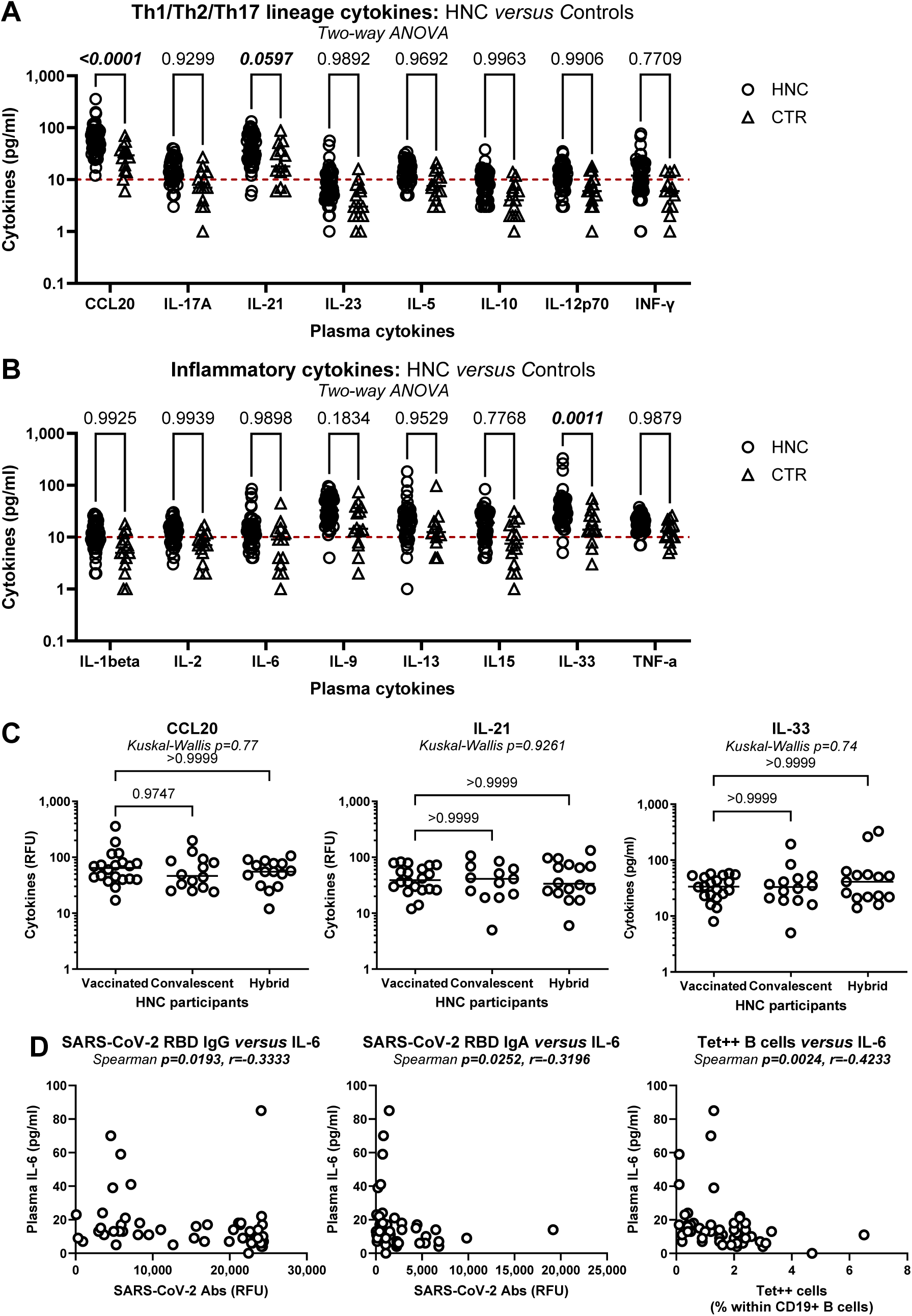
Plasma levels of Th1/Th2/Th17 and inflammatory cytokines in HNC participants with distinct immunization status. Plasma levels of 25 cytokines were quantified using the Luminex xMAP-based multiplex assay. (**A-B**) The depicted Th1/Th2/Th17 (**A**) and inflammatory cytokines (**B**) were present at detectable levels in plasma of HNC and control participants. Two-way ANOVA test followed by Sidak’s multiple comparisons determined differences between groups, with p-values indicated on the graphs. (**C**) Shown are differences in CCL20, IL-21, and IL-33 levels between HNC participants classified based on the immunization status in vaccinated, convalescents and hybrid, with Kruskall-Wallis and Dunn’s multiple comparisons p-values indicated on the graphs. (D) Spearman p and r values are indicated for the correlations between plasma levels of IL-6 and plasma levels of SARS-CoV-2 RBD IgG (**right panel**) and IgA (**middle panel**) Abs, as well as the frequency of Tet++ B-cells (**left panel**) in HNC participants.

### Linear regression to identify correlates of SARS-CoV-2-specific humoral immunity outcomes

Plasma levels of RBD Abs with IgG or IgA isotypes may represent surrogate markers of SARS-CoV-2 neutralization potential, while the frequency of Tet^++^ B-cells, likely producing these neutralizing Abs, reflects the persistence of humoral immunity upon vaccination and/or natural infection. To identify predictors of these three major humoral immunity outcomes (*i.e.,* RBD IgG, RBD IgA, Tet^++^ B-cells) among the panoply of immunological measurements realized in this study, a linear regression model was used. The results, including the regression coefficient, p-values and adjusted p-values (Adj. p) are presented in Supplemental Tables 1-4. A negative/positive coefficient indicated a negative/positive correlation. In addition to the crude model, adjustments for numerical (*i.e.,* age, BMI and TSI) and/or categorical (*i.e.,* sex, smoking, alcohol, toxic environment, diabetes) variables were performed (Supplemental Tables 1-4).

For SARS-CoV-2 Abs measurements depicted in Figure 1, levels of S1, S2, and NC IgG Abs positively correlated with RBD IgG Abs levels, while levels of S1 and S2 IgG and IgA Abs levels positively correlated with RBD IgA Abs levels, with statistical significance (Adj. p<0.05) maintained upon adjustment for numerical and categorical variables (Supplemental Table 1). For B-cell measurements (Figures 2-3), the frequencies of total B-cells with a memory phenotype (CD27^+^) and IgA isotype negatively correlated with RBD IgG Abs levels, while the frequency of total and IgM^+^Tet^++^ B-cells negatively correlated with the frequency of total Tet^++^ B-cells (Supplemental Table 2). In contrast, the frequency of IgA^+^Tet^++^ B-cells positively correlated with the frequency of total Tet^++^ B-cells (Supplemental Table 2). For readouts on CD4^+^ and CD8^+^ T-cell proliferation *in vitro* (Figure 4), none significantly predicted the three major SARS-CoV-2 immunity outcomes (Supplemental Table 3). Finally, for read-outs on plasma cytokines (Figure 6), there was a tendency for a negative correlation of IL-6 levels with plasma RBD IgG and IgA levels, and the frequency of Tet^++^ B-cells, reaching statistical significance only for p-values, upon adjustment for numerical and/or categorical variables (Supplemental Table 4). Additionally, plasma IL-13 levels were also negatively correlated with RBD IgA levels and with the frequency of Tet^++^ B-cells, with p-values reaching statistical significance mainly upon adjustment for numerical and categorical variables (Supplemental Table 4).

In conclusion, we identified multiple positive and negative predictors of three major SARS-CoV-2 immunity outcomes, as resumed in Table 2.

**Table 2.**
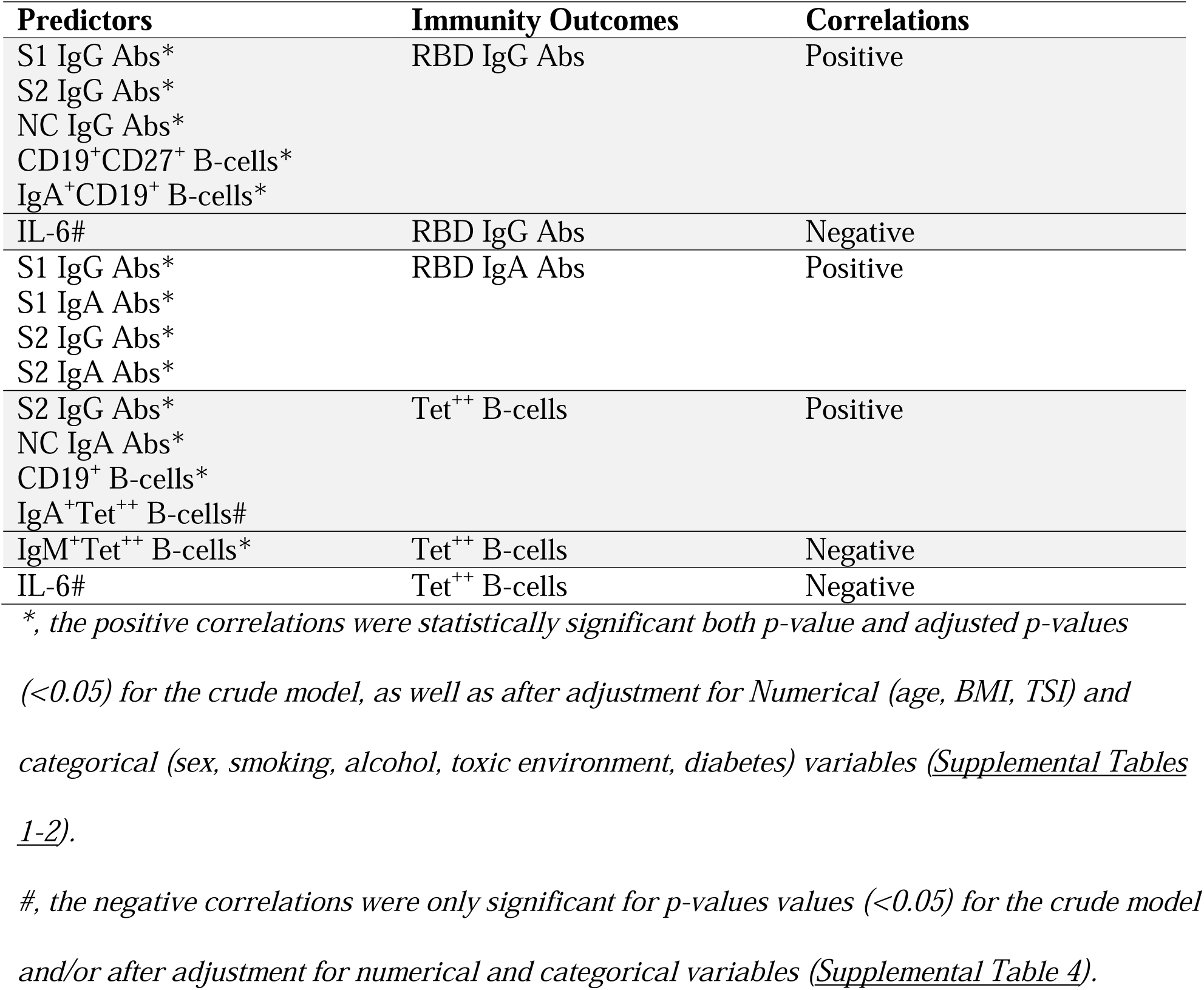
Humoral predictors of the main SARS-CoV-2 immunity outcomes revealed in the linear regression model.

### Long-term persistence of Tet^++^ B-cells in HNC patients

To explore the duration of SARS-CoV-2-specific B-cell immunity in HNC patients, the frequency of Tet^++^ B-cells was monitored in a subgroup of HNC participants (n=25) at Visit 1 and at Visit 2 (median TSI 117 and 341 days, respectively) (Figure 7A). In terms of total CD19^+^ B-cells, a decrease in their frequency was observed at Visit 2 *versus* 1, with no changes in the expression of the memory marker CD27, but with an increased frequency of the IgM isotype (Figure 7B-C). Variations in the frequency of Tet^++^ B-cells from Visit 1 to 2 were heterogeneous among participants, with overall no statistically significant differences between the two Visits (Figure 7D, <u>left panel</u>). The immunization status was not associated with a particular trend in the persistence of Tet^++^ B-cells (<u>data not shown</u>). However, a decreased frequency of CD27^+^ Tet^++^ B-cells was observed at Visit 2 *versus* 1 (Figure 7D, <u>right panel</u>), without changes in the isotype of Tet^++^ B-cells (Figure 7E). Finally, in this group of HNC participants the TSI negatively correlated with the frequency of Tet^++^ B-cells at Visit 1 but not at Visit 2 (Figure 7F). Finally, the stratification by immunization status revealed no significant differences in the frequency of Tet^++^ B-cells between Visit 1 and Visit 2 among the HNC groups (Figure 7G). All together, these results indicate the long-term persistence of circulating Tet^++^ B-cells in HNC patients, although with a reduction in the memory phenotype, up to day 717 post-immunization.

**Figure 7:**
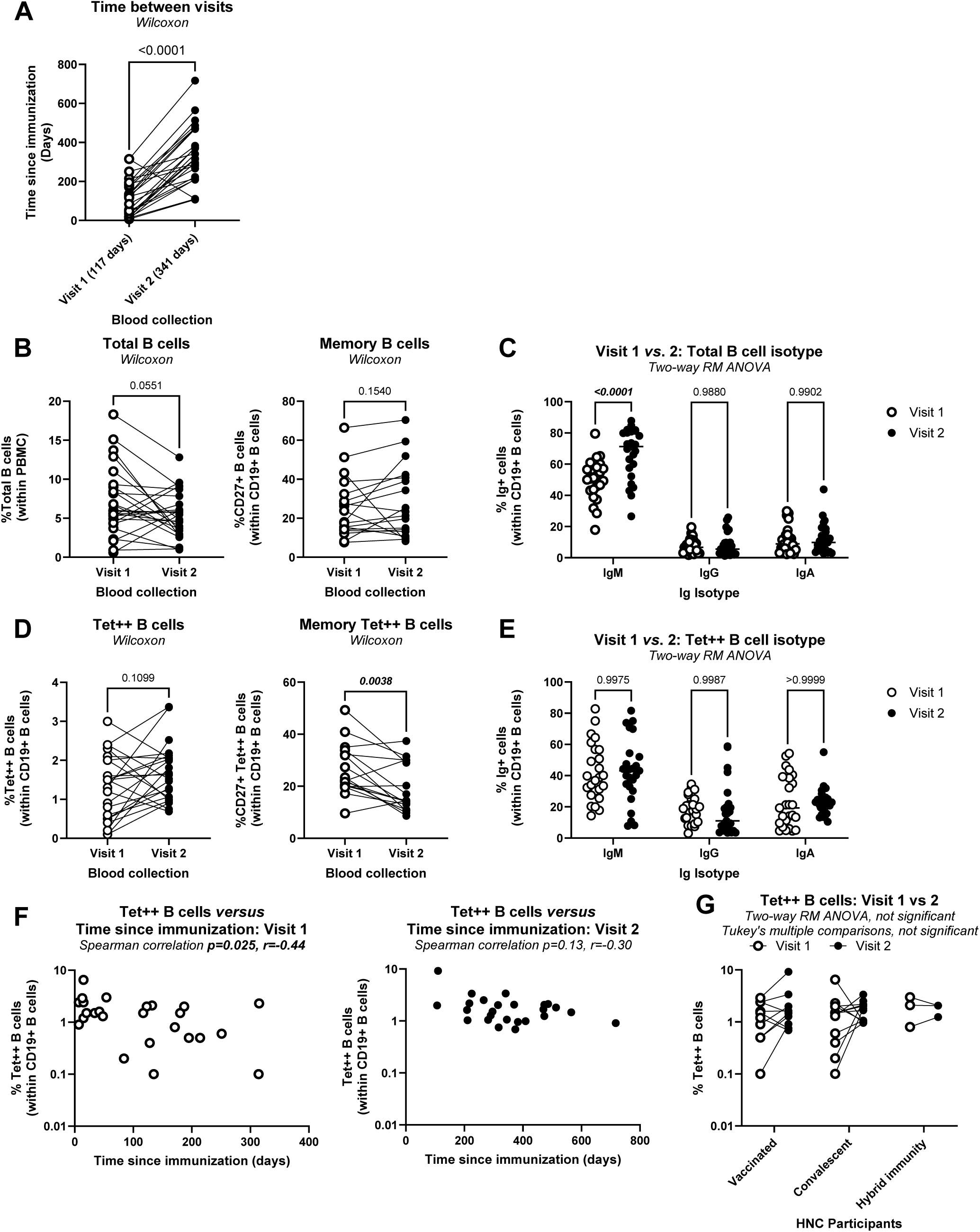
Long-term persistence of SARS-CoV-2 RBD-specific B-cells in the blood of HNC participants. The frequency of total and Tet^++^ B-cells with a memory CD27^+^ phenotype and IgM, IgG, or IgA isotypes, identified by flow cytometry as detailed in Figure 2 <u>legend</u>, was analyzed at two distinct visits in a group of n=25 HNC participants. (**A**) Shown is the time since immunization at visit 1 and visit 2 (median time since immunization 117 and 341 days, respectively; as indicated in parenthesis). (**B-C**) Shown are the frequencies of total (**B, left panel**) and memory (CD27^+^) (**B, right panel**) CD19^+^ B-cells, as well as their isotype (**C**), at visit 1 compared to visit 2. (**D-E**) Shown are the frequencies of total (**D, left panel**) and memory (CD27^+^) (**D, right panel**) Tet^++^ CD19^+^ B-cells, as well as their isotype (**E**), were compared between visit 1 and 2. Wilcoxon p-values (**A, B and D**) and Two-way ANOVA test followed by Sidak’s multiple comparisons (**C and E**) determined differences between groups. **(F)** Shown is the correlation between the frequency of Tet^++^ B-cells and the time since immunization at Visit 1 (**left panel**) and Visit 2 (**right panel**), with Spearman correlation p and r values indicated on the graphs. **(G)** Comparison of the frequency of Tet^++^ B-cells between Visit 1 and Visit 2 among the HNC participants.

## DISCUSSION

In this study, we investigated the amplitude and the duration of natural and vaccine-induced SARS-CoV-2 immunity in a cohort of people with HNC (n=49) admitted for oncologic treatment at Coltea Hospital, Bucharest, Romania. Plasma samples were used for the measurement of SARS-CoV-2 Spike S1, S2 and RBD, and NC Abs, as well as of cytokines, while PBMCs were used for the monitoring of RBD-specific B-cells (Tet^++^) *ex vivo*, and the proliferation of B-cells and CD4^+^ and CD8^+^ T-cells in response to SARS-CoV-2 Spike and NC *in vitro*. Our results reveal the ability of HNC to develop efficient SARS-CoV-2 immunity in response to vaccination and/or natural infection, responses that persisted long-term (median TSI: 341 days). Our results also confirm the value of “*hybrid immunity*” in HNC patients. These results have clinical relevance for the management of HNC patients in the context of the persistent circulation of SARS-CoV-2 VOCs.

The highly pathogenic SARS-CoV-2 variants alpha, beta, gamma, and delta, originally identified in UK, South Africa, Brazil, and India, respectively, emerged in September 2020 from the ancestral virus that underwent multiple escape mutations^101,102^. The less pathogenic omicron variants, initially detected in South Africa, distinguished from the other VOCs by their capacity to infect immunized individuals^102,103^. Enrollment of participants in our study at Visit 1 started in August 2021 and finalized in February 2022, with the majority of participants recruited in August-December 2021, when the SARS-CoV-2 delta variant was predominant in Romania.

In our cohort, unvaccinated HNC participants infected with SARS-CoV-2 developed only mild to moderate symptoms. This could be explained by the fact that HNC patients detected positive for SARS-CoV-2 by RT-PCR prior hospital admission received prompt treatment using the available antiviral and anti-inflammatory drugs and were followed-up remotely by the physicians until the resolution of symptoms. Another reason might be the natural resistance to disease, consistent with the reported decreased expression of SARS-CoV-2 receptors/co-receptors in tumor tissues of HNC patients^104–108^. In line with this possibility, a very recent study provides evidence than HNC patients have a genetic predisposition for a mild form of COVID-19^109^. Moreover, a very recent Mendelian randomization study found no link between genes identified in *Gene-Wide Association Studies* (GWAS) and susceptibility to COVID-19 and HNC, and suggested a positive effect of viral infections on cancer outcome *via* mechanisms depended on the type I interferon alpha receptor 2 (IFNAR2)^110^. Also, it is noteworthy that one study performed in Denmark concluded that HNC patients are not at high-risk of acquiring SARS-CoV-2 infection^111^.

Our study demonstrated that levels SARS-CoV-2 Abs and the frequencies of Tet^++^ B-cells did not differ between HNC and control groups, with the two groups being similar in age, TSI and BMI. One way to distinguish between convalescent and vaccinated individuals is the detection of humoral/cellular immune responses against the NC, responses induced only upon natural infection^32,33,37^. Consistently, in our cohort, the detection of NC Abs, as well as B-cells and CD4^+^ and CD8^+^ T-cells proliferating in response to recombinant NC proteins was observed only in convalescent and hybrid immunity participants. It is to be noticed that the highest levels of SARS-CoV-2 Abs, including those directed against Spike S1, S2, and RBD proteins of IgG and IgA isotypes, were detected in individuals that were exposed to natural infection before or after vaccination, a phenomenon called “*hybrid immunity*”^112–116^. Multiple studies demonstrated the superiority of hybrid immunity in terms of amplitude, breath, and duration of protective SARS-CoV-2-specific immune responses^112–116^. Similar to plasma levels of SARS-CoV-2 Abs, the frequency of Tet^++^ B-cells and the proliferation (CFSE^low^) of B-cells and CD4^+^/CD8^+^ T-cells in response to recombinant Spike proteins, were the most robust in HNC patients with hybrid immunity compared to vaccinated and convalescent participants. Compared to total B-cells, Tet^++^ B-cells were enriched in IgG and IgA isotypes, although the IgM isotype remained predominant, with no differences in the Ig isotypes between vaccinated, convalescent and hybrid immunity in the HNC group. Of note, there was a significant positive correlation between the frequency of Tet^++^ B-cells and plasma levels RBD Abs of both IgG and IgA isotypes, suggesting that circulating Tet^++^ B-cells represent the source of these RBD Abs. Also, the proliferation of CD4^+^ T-cells in response to Spike proteins was positively correlated with the frequency of Tet^++^ B-cells, in line with the importance of CD4^+^ T-cells (*e.g.,* follicular helper) in sustaining B-cell proliferation^22,23^.

Protection against infection is insured by Abs able to neutralize SARS-CoV-2 entry into host cells^13,15^. SARS-CoV-2 Abs directed against RBD typically exhibit neutralizing capacity^117^. Therefore, among the multitude of immune parameters monitored in this study, plasma levels of RBD Abs of IgG and IgA isotypes, and the frequency of circulating RBD-specific Tet^++^ B-cells, were considered the major immunity outcomes. A linear regression model was used to identify immunological predictors of these immunity outcomes. For the <u>RBD IgG Ab outcome</u>, the positive predictors identified included the S1, S2 and NC Abs of IgG isotype only, and the frequency of CD19^+^CD27^+^ B-cells and CD19^+^ B-cells with an IgA isotype. For the <u>RBD IgA Ab</u> <u>outcome</u>, the positive predictors identified included S1 and S2 Abs of both IgG and IgA isotypes. Finally, for the Tet^++^ B-cell outcome, the positive predictors identified included S2 IgG and NC IgA Abs, as well as the frequency of CD19^+^ B-cells and that of Tet^++^ B-cells with an IgA isotype, while the frequency of IgM^+^Tet^++^ B-cells was a negative predictor. These predictors remained in majority significant upon adjustments based on numerical (age, BMI, TSI) and categorical (sex, smoking, alcohol, toxic environment, diabetes) cofounding factors.

Among the panel of 25 cytokines quantified in the plasma, a significant increase in HNC *versus* control groups was only observed for CCL20, IL-21 and IL-33. Of note, CCL20 was shown to be upregulated in HNCC patients, contributing to tumor progression^118^, IL-21 is produced by follicular helper CD4^+^ T-cells that sustain B-cell functions^119^, while IL-33 levels are associated which exacerbated pulmonary inflammation in COVID-19^120^. Among HNC participants, CCL20, IL-21 and IL-33 levels did not vary based on the immunization status. However, Spearman correlations revealed that plasma levels of IL-6, and at a lower extend IL-10 and IL-13, were negatively correlated with plasma levels of RBD IgG and IgA Abs, and the frequency of Tet^++^ B-cells. In addition, plasma levels of IL-6 were also identified as moderate negative predictors for the frequency of Tet^++^ B-cells in the linear regression model, with and without adjustment for numerical and categorical confounding factors. This is consistent with the knowledge that IL-6 is a marker of SARS-CoV-2 infection severity, and that Tocilizumab, an anti-IL-6 Abs, is used for the treatment for COVID-19 pneumonia^55,121–123^.

Finally, the longitudinal follow-up in a fraction of n=25 HNC participants revealed the persistence of Tet^++^ B-cells at similar frequencies between two Visits, at a median TSI of 117 (Visit 1) and 341 days (Visit 2). Similar to findings at Visit 1, a major fraction of Tet^++^ B-cells exhibited an IgM isotype, while minor fractions presented IgG and IgA isotypes at Visit 2. However, a decreased frequency of Tet^++^ B-cells with and CD27^+^ memory phenotype was observed at Visit 2 *versus* Visit 1, indicating either a decline in the survival capacity of Tet^++^ B-cells or a more recent activation of these cells, considering the strong likelihood of SARS-CoV-2 re exposure and/or reinfection in 2022-2023. Nevertheless, the persistence of Tet^++^ B-cells up to 717 days post-immunization (median TSI: 341 days) in this cohort of HNC patients, points to a robust humoral immunity in HCN individuals, similar to the general population^32,124^. This aligns with findings from other studies on oncologic patients^131, 132^, reinforcing the evidence that HNC patients possess the immunological competence to develop an effective immune response against SARS-CoV-2 infection.

This study has several limitations we acknowledge here. The control group (n=14) was small in size and younger compared to the HNC group (n=49). Due to sample size limitations, we were unable to stratify HNC participants *per* cancer pathology and type of vaccines, aspects that can influence the immune responses^125^. Nevertheless, the stratification of HNC patients based on the immunization status was possible and confirmed the superiority of *hybrid immunity*, as in previous studies^112–116^. The underrepresentation of women in our HNC cohort is attributed to the highest prevalence of HNC in males^88–91^. Our immune monitoring studies were confined solely to analyses of peripheral blood. Finally, we acknowledge the importance of investigating respiratory mucosal immunity, which are key for SARS-CoV-2 transmission^126^ and immunity^127,128^.

In conclusion, this study provides an original analysis of the breadth and duration of humoral and cellular immunity to SARS-CoV-2 among a cohort of 49 HNC patients receiving oncologic treatment from Bucharest, Romania. Briefly, the key findings include: ***i)*** similarities in three major immunological outcomes (*i.e.,* plasma levels of RBD IgG and IgA Abs, and the frequency of Tet^++^ B-cells) between HNC and control groups; ***ii)*** the strongest immunological responses were observed in participants with hybrid immunity; ***iii)*** the persistence of Tet^++^ B-cells in peripheral blood for up to 717 days post-immunization (median TSI: 341 days), suggesting durable protective immunity in HNC patients. Longitudinal follow-up studies are required to validate these three major immunological outcomes as predictors of long-term protection against SARS-CoV-2 re-infection in the general population *versus* people with HNC. Together, these investigations open the path for further research into the relationship between antiviral and antitumor immunity in specific oncological populations.

## Contributors

LM contributed to study design, performed the majority of experiments, analyzed data, prepared figures, and wrote the manuscript. AE obtained ethical approval for the study, recruited participants, collected biological samples, and provided access to clinical information. NMA performed experiments, and contributed to figure preparation. MN, DAC, MP, and ED contributed to blood processing and plasma/PBMC sample storage, performed preliminary experiments, and contributed to figure preparation. EC and VR provided protocols and expertise with multiplex assays for the quantification of Abs and cytokines. BG and AH provided protocols and expertise with polychromatic flow cytometry experiments. GG and IB provided protocols and reagents for preliminary quantification of Abs, and provided student supervision. AMF performed statistical analysis and prepared tables. VL contributed to study design, obtained ethical approval, and contributed to manuscript writing. CC contributed to study design, provided protocols, supervised staff, secured funding for salary support and operation, and contributed to manuscript writing. RG designed the study, obtained ethical approvals, recruited participants, provided access to biological samples and clinical information, and contributed to manuscript writing. PA conceived the research study, provided protocols, optimized and performed experiments, supervised students, analyzed data, prepared figures, performed statistical analysis, and wrote the manuscript. All authors revised and approved the final version of the manuscript and had access to the raw data.

## Acknowledgments

The authors acknowledge the key contributions of all study participants for their gift of blood samples essential for this study. PA acknowledges salary support from the Université de Montréal for sabbatical year (07/2021-06/2022) and a salary complement from the University of Bucharest. LM acknowledges funding from the Executive Agency for Higher Education, Research, Development and Innovation Funding (UEFISCDI, Romania) grant number PN-III-P2-2.1-PED-2021-2115 (620 PED). NMA, MN and DAC received Excellence M.Sc. Fellowships from the University of Bucharest, Romania.

## Data sharing statement

All data generated are included in the Figures, Tables and Supplemental Figures, Tables and Files. Access to primary data is available upon request to the lead author: petronela.ancutamontreal.ca

