## Supplemental Figures 1-3 for "Unveiling Humoral and Cellular Immune Responses to SARS-CoV-2 in Head and Neck Cancer: A Comparative Study of Vaccination and Natural Infection in Romania"

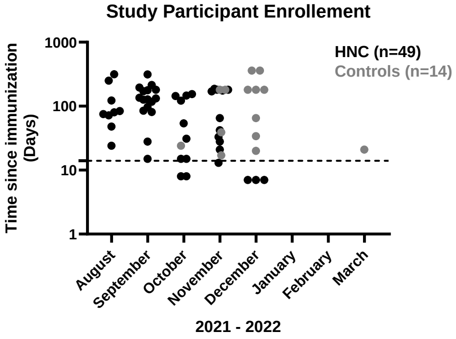


**Supplemental Figure 1 (related to Figures 1-7): Time of study participant recruitment.** The study cohort included n=49 HNC and n=14 control participants recruited at Coltea Hospital (Bucharest, Romania). Plasma and PBMCs specimens were collected at one (n=49 HNC participants) or two (n=25 HNC participants) subsequent visits between August 2021 and March 2022.

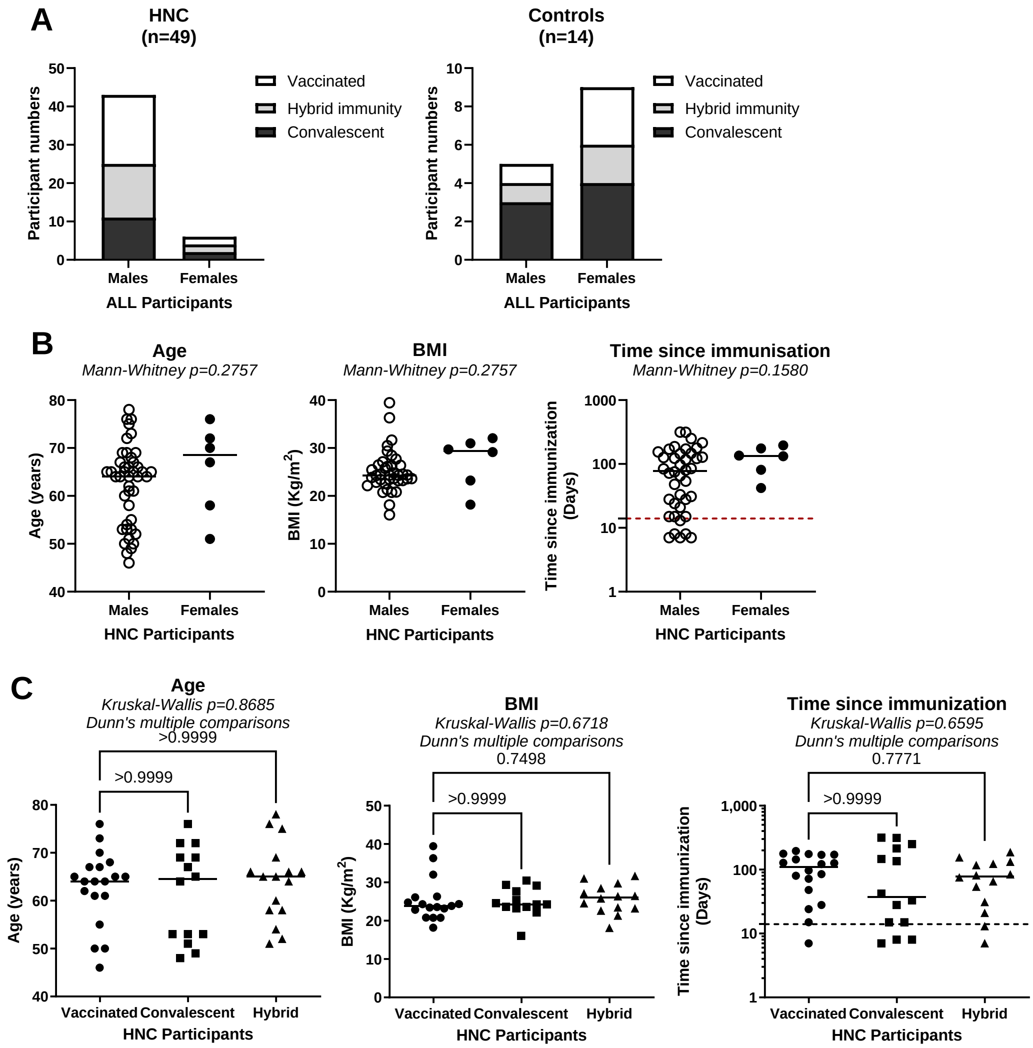


**Supplemental Figure 2 (related to Figures 1-7): Sex, age, body mass index and time since immunization of study participants.** The HNC (n=49) and control (n=14) study participants were analyzed based on age, body mass index (BMI), time since immunization, and immunization status characteristics at baseline (Visit 1). (**A**) Shown is the distribution of the three groups: vaccinated, convalescents and hybrid immunity among the HNC (**A, left panel**) and control (**A, right panel**) participants separated by sex. (**B**) Show are differences in age, BMI and time since immunization between male and female HNC participants. Mann-Whitney p-values are indicated on the graphs. (**C**) Shown are differences in age, BMI and time since immunization between HNC the three groups of HNC participants (vaccinated, convalescents and hybrid immunity). Kruskal Wallis with subsequent Dunn’s multiple comparisons test p-values are depicted on the graphs.

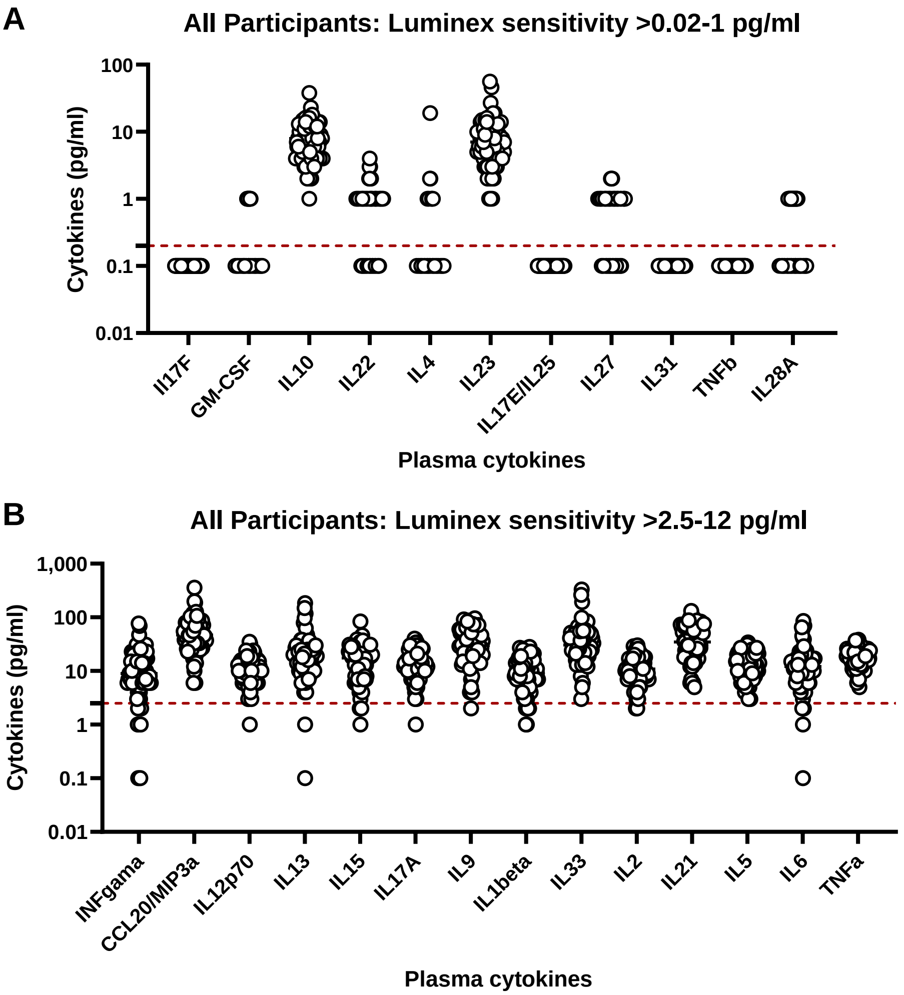


**Supplemental Figure 3 (related to Figure 6): Detection of plasma cytokines in HNC participants**. Plasma levels of 25 cytokines were quantified using the Luminex xMAP-based multiplex assay. Shown are levels of cycokines present in the plasma classified based on their relatively low (**A;** sensitivity threshold 0.02-1 pg/ml) and high (**B;** sensitivity threshold 2.5-12 pg/ml) levels of expression. The red dashed line indicates the threshold of detection.
